# Behavioral, Biochemical and Histopathological effects of Standardised Pomegranate extract with Vinpocetine, Propolis or Cocoa in a rat model of Parkinson’s disease

**DOI:** 10.1101/2020.06.03.131615

**Authors:** Azza A. Ali, Mona M. Kamal, Mona G. Khalil, Shimaa A. Ali, Hemat A. Elariny, Amany Bekhit, Ahmed Wahid

**Author notes:** Corresponding author Ahmed Wahid, Department of Pharmaceutical Biochemistry, Faculty of Pharmacy, Alexandria University, Alexandria, Egypt.

## Abstract

**Introduction:** Parkinsonism is a neurodegenerative disorder. Pomegranate (POM) has been previously shown to have a dopaminergic neuroprotective effect against Parkinsonism.

**Objective:** The aim of the current study is to compare the efficacy of POM, vinpocetine, Propolis, Cocoa or L-dopa using RT-induced Parkinsonism rat model.

**Methods:** Rats were divided into seven groups; one normal and five RT model groups. One of the RT (2.5 mg/kg sc) groups served as non-treated parkinsonism model whereas the others were treated with either L-dopa (10 mg/kg PO) or with POM (150 mg/kg PO) together with each of the following; vinpocetine (VIN) (20 mg/kg PO), Propolis (300 mg/kg PO), Cocoa (24 mg/kg PO). Motor and cognitive performances were examined using three tests (catalepsy, open-field, Y-maze). Striatal dopamine, norepinephrine, serotonin, acetylcholinesterase, GABA, Glutamate, GSK 3B, BDNF levels were assessed as well as MDA, SOD, TAC, IL-1β, TNF-α, iNOs and caspase-3. Also, histopathological examinations of different brain regions were determined.

**Results:** Treatment with L-dopa alone or with all POM combination groups alleviated the deficits in locomotor activities, cognition, monoamine levels, acetylcholinesterase activity, oxidative stress, and inflammatory markers as well as caspase-3 expression induced by RT.

**Conclusion:** Combinations of POM with each of VIN, Propolis or Cocoa have a promising disease-modifying antiparkinsonian therapy even without being given as an adjuvant to L-dopa.

## Introduction

Neurodegenerative diseases (NDD) were reported to affect millions of people each year all over the world. But yet the complex multifactorial mechanisms of parkinsonism are not fully understood so far. It has been proposed previously that Manganese can cause severe neurological damage [1]. Parkinsonism is defined as the presence of bradykinesia, rest tremor and rigidity. The exact etiology of Parkinsonism remains elusive. Age is the most significant risk factor for the development of Parkinsonism. In addition to that, complex interactions between environmental and genetic factors can be a predisposing factor to Parkinsonism.

It was previously demonstrated that the progression of neurodegeneration such as the case in parkinsonism is associated with decreased antioxidant levels and increased oxidative damage to proteins, DNA and lipids [2]. Unquestionably the injurious effects of the inflammatory response are associated with augmentation of ROS and oxidative damage that was assessed by inhibition of defense mechanisms such as superoxide dismutase (SOD), glutathione (GSH). Oxidative stress can then result in induction of the gene expression of a battery of distinct pro-inflammatory mediators such as, TNF-α, interleukin-1-beta (IL-1β).

Natural products are needed as a substitute for chemical therapeutics for their safe toxicity profiles. There is ample evidence that oxidative stress and inflammation are involved in the pathogenesis of parkinson’s disease (PD), therefore, the use of natural agents modulating both oxidative stress and inflammation have been proposed as a mainstream choice to counteract PD.

This study aims to investigate the efficacy of POM together with each of VIN, Propolis, Cocoa or L-dopa using rotenone-induced PD rat model.

## Materials and Methods

### Animals

Adult male albino rats, weighing approximately 300-340 g at the beginning of the experiment, were obtained from the Nile Co. for Pharmaceuticals and Chemical Industries, Cairo, Egypt. Animals were kept under the same adequate environmental conditions and provided with their daily dietary requirements consisting of standard diet pellets (El-Nasr Chemical Co., Abu Zaabal, Cairo, Egypt) and water was given ad-libitum. Rats were housed in stainless-steel cages, three per cage and kept at the animal house (at the facility of Faculty of Pharmacy, Al-Azhar University “girls”). Animal experiments followed the national institute of health guidelines for the care and use of laboratory animals (NIH publications No. 8023, revised 1978). Animal experiments were usually carried out at a fixed time around 9 am-2 pm. All experimental procedures of the study were approved by the animal care and use committee of the Faculty of Pharmacy, Al-Azhar University (ethical approval number 212).

### Chemicals and reagents

All of the chemicals and reagents used in the present study were of high analytical Grade. Chemicals were purchased from Sigma-Aldrich Co. Rotenone was obtained as a white to dark beige powder. It was suspended in sunflower oil to a concentration of 2.5 mg/ml. Whereas, Pomegranate extract standardized to 40% ellagic acid polyphenol was purchased as a grey-brown powder from Holland &Barret, UK. It was dissolved in 10 % Tween 80 saline solution.

### Experimental design

Rats were divided into seven groups and treated with different drugs for 30 days as follows:

**Group I**: Normal healthy control group. **Group II**: RT (2.5 mg/kg sc) served as non-treated Parkinsonism model. **Group III**: RT (2.5 mg/kg sc) and L-dopa (10 mg/kg PO) [3]. **Group IV**: RT (2.5 mg/kg sc), POM (150 mg/kg PO) [4] and VIN. (20 mg/kg PO) [5]. **Group V**: RT (2.5 mg/kg sc), POM (150 mg/kg PO) and Cocoa (24 mg/kg PO) [6]. **Group VI**: RT (2.5 mg/kg sc), POM (150 mg/kg PO) and Propolis (300 mg/kg PO) [7]. **Group VII**: RT (2.5 mg/kg sc), POM (150 mg/kg PO) and L-dopa (10 mg/kg PO).

The following investigations were then conducted:

### Behavioral Assessment

Behavioral investigations such as Open-Field test (locomotors activity, emotionality and exploratory behavior assessment), Catalepsy tests such as Grid test and swimming test, in addition to Y-maze test (Assessment of spatial working memory) were conducted as done previously [8,9]. After the last behavioral test, rats were sacrificed by decapitation and the brains removed for postmortem biochemical and histological assessments. Diethyl ether (Merck) was used as an anesthetic agent during obtaining of the brain from rats.

### Biochemical investigations

The activity of BDNF, IL1β, TNF-α, GABA, GSK-3B,MDA, TAC, SOD and Caspase-3, iNOs, Glutamate in striatal tissue homogenate as well as monoamine contents (DA, NE, and 5-HT) in frontal cortex and striatum were carried out using commercially available test ELISA kits supplied by Biodiagnostic, Giza, Egypt following the manufacturers’ instructions. Acetylcholinesterase activity was assessed using a kit obtained from Sigma-Aldrich Co., St Louis, MO, USA.

### Histopathological Examination of Different Brain Regions

Brain specimens were fixed in 10% formalin for 24 h then washed with tap water and done as previously described [10].

### Statistical Analysis

Data were expressed as the mean ± SEM and statistical analysis was carried out by one way ANOVA followed by Tukey multiple comparisons test to calculate the significance of the difference between treatments. Values of p < 0.05 were considered significant. All statistical analyses were performed and graphs were sketched using GraphPad Prism (ISI, USA) software (version 5) computer program.

## Results

In the present work, the concurrent administration of L-dopa alone or with all POM combinations could reverse the deficits of Rotenone in rat animal model of Parkinsonism. This observation was confirmed via conducting three different types of experiments: Behavioural, Biochemical and histopathological investigations.

### The Behavioral efficacies of Pomegranate with each of vinpocetine, propolis, cocoa or L-dopa against the development of parkinsonism in a rat animal model

To investigate the neuroprotective potential of POM and other compounds employed in this study, we conducted several behavioral analyses such as open field, catalepsy, and Y-maze tests.

### Open-field tests

The open-field test is a mild stressful condition. It is a general test for locomotor activity in rodents. To investigate the neuroprotection effect of POM, Rotenone was administered as 2.5 mg/kg sc for 30 consecutive days. Rotenone group showed a significant decrease in locomotion, which was shown in a decreased number of ambulation, rearing and grooming frequencies. POM with different combinations employed in the current study significantly enhanced the motor activity of the rats in the open field test as compared to rotenone group. The number of ambulations for the groups: control, Rotenone, Rotenone and L-dopa, Rotenone, POM, and VIN, Rotenone, POM, and cocoa, Rotenone, POM, and propolis, Rotenone, POM and L-dopa, were respectively as follows: 31.83± 6.555, 9.667±2.338, 23.67±5.785, 25.67±5.854, 25.83±5.037, 28.00±6.753, 27.83±3.251) (see figure 1).

**Figure 1:**
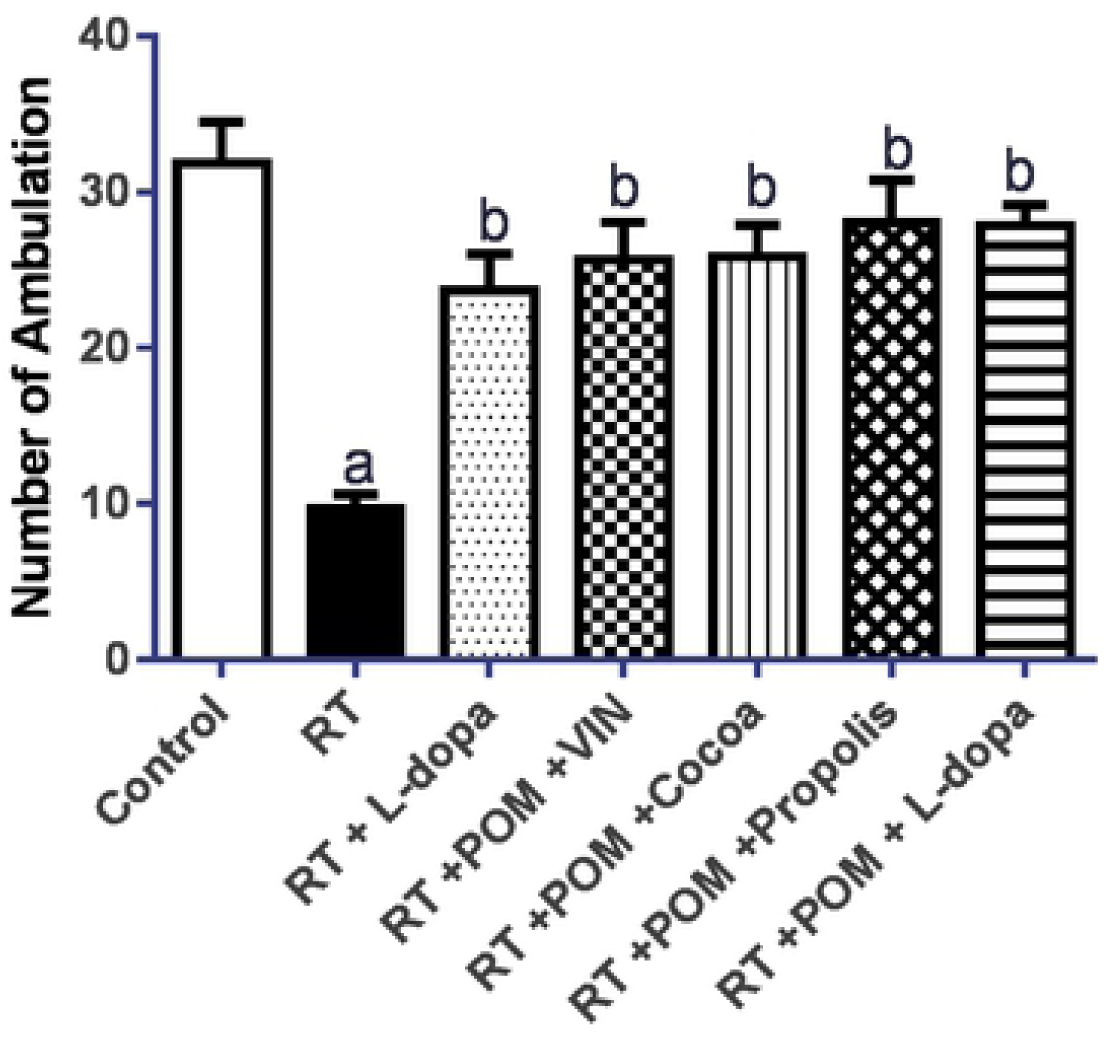
Neurobehavioural assessment: Ambulation frequency in an open field test: a, b denote significant difference from control, and Rotenone groups respectively at P <0.05.

Similarly, Rotenone has the capability to minimize significantly the rearing frequency (p < 0.05), in comparison to the respective control group. The number of rearings for the groups: control, Rotenone, Rotenone and L-dopa, Rotenone, POM, and VIN, Rotenone, POM, and cocoa, Rotenone, POM, and propolis, Rotenone, POM and L-dopa, were respectively as following: 19.33±3.724, 4.333±1.033, 10.00±2.449, 12.33±3.011, 15.50±3.271, 16.00±3.521, 15.67±2.875) (see figure 2).

**Figure 2:**
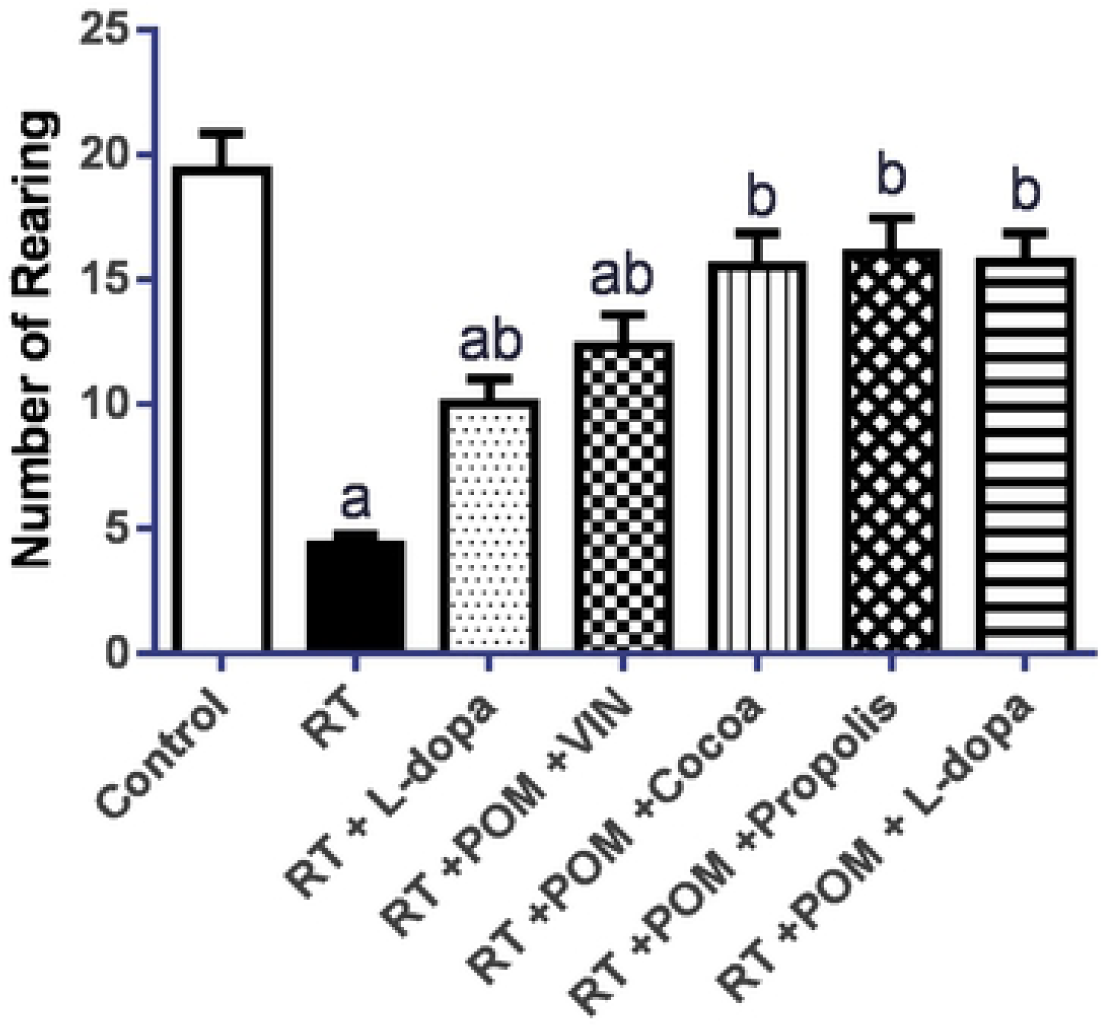
Neurobehavioural assessment: Rearing frequency in an open field test: a, b denote significant difference from control, and Rotenone groups respectively at P <0.05.

Furthermore, the number of groomings showed the same pattern of ambulations and rearings. The number of groomings for the groups: control, Rotenone, Rotenone and L-dopa, Rotenone, POM, and VIN, Rotenone, POM, and cocoa, Rotenone, POM, and propolis, Rotenone, POM and L-dopa were respectively as follows: 25.20± 3.162, 5.830±1.140, 16.80±1.581, 19.50±1.581, 23.00±2.025, 23.00±1.703, 23.50±1.703 seconds) (see figures 1-3).

**Figure 3:**
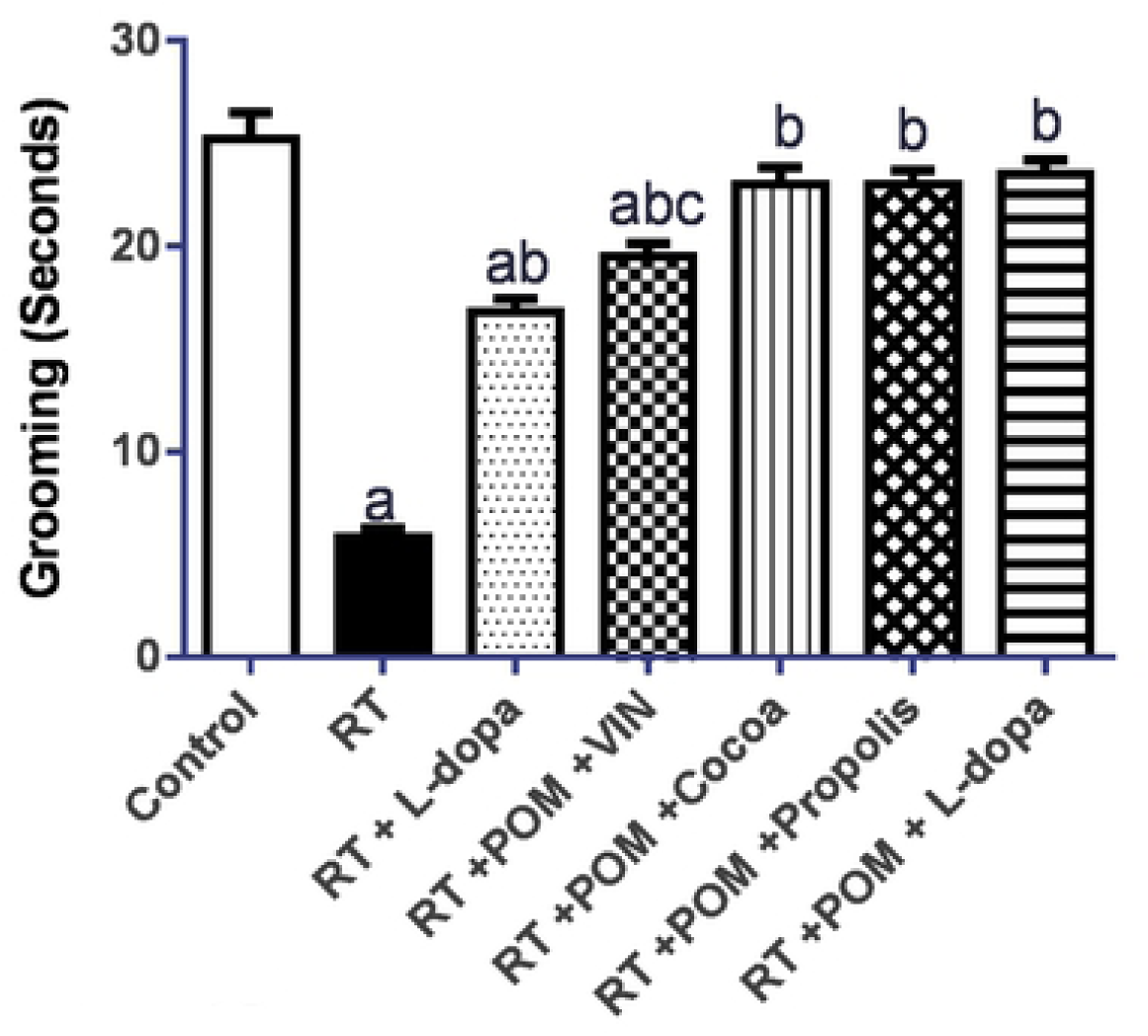
Neurobehavioural assessment: Grooming frequency in an open field test: a, b, c denote significantly different from control, Rotenone, and Rotenone/L-DOPA groups respectively at P <0.05.

### Catalepsy test

Catalepsy test is an index for the presence or absence of parkinsonian-like symptoms such as “bradykinesia, akinesia, and rigidity”. Male rats were tested on the grid test and used as an index of catalepsy or sensorimotor deficit. Rotenone showed the highest moving latency, whereas L-dopa or POM alone and/or in combination with VIN, Cocoa, and Propolis showed significant improvements in comparison to the Rotenone group (P<0.05). Moving latency for the groups: control, rotenone, rotenone and L-dopa, rotenone, POM, and VIN, rotenone, POM, and cocoa, rotenone, POM, and propolis, rotenone, POM and L-dopa, were respectively as follows: 1.083±0.2041, 3.000±0.6325, 2.083±0.4916, 1.083±0.2041, 1.283±0.3189, 1.267±0.2944, 1.250±0.2739 seconds) (see figure 4).

**Figure 4:**
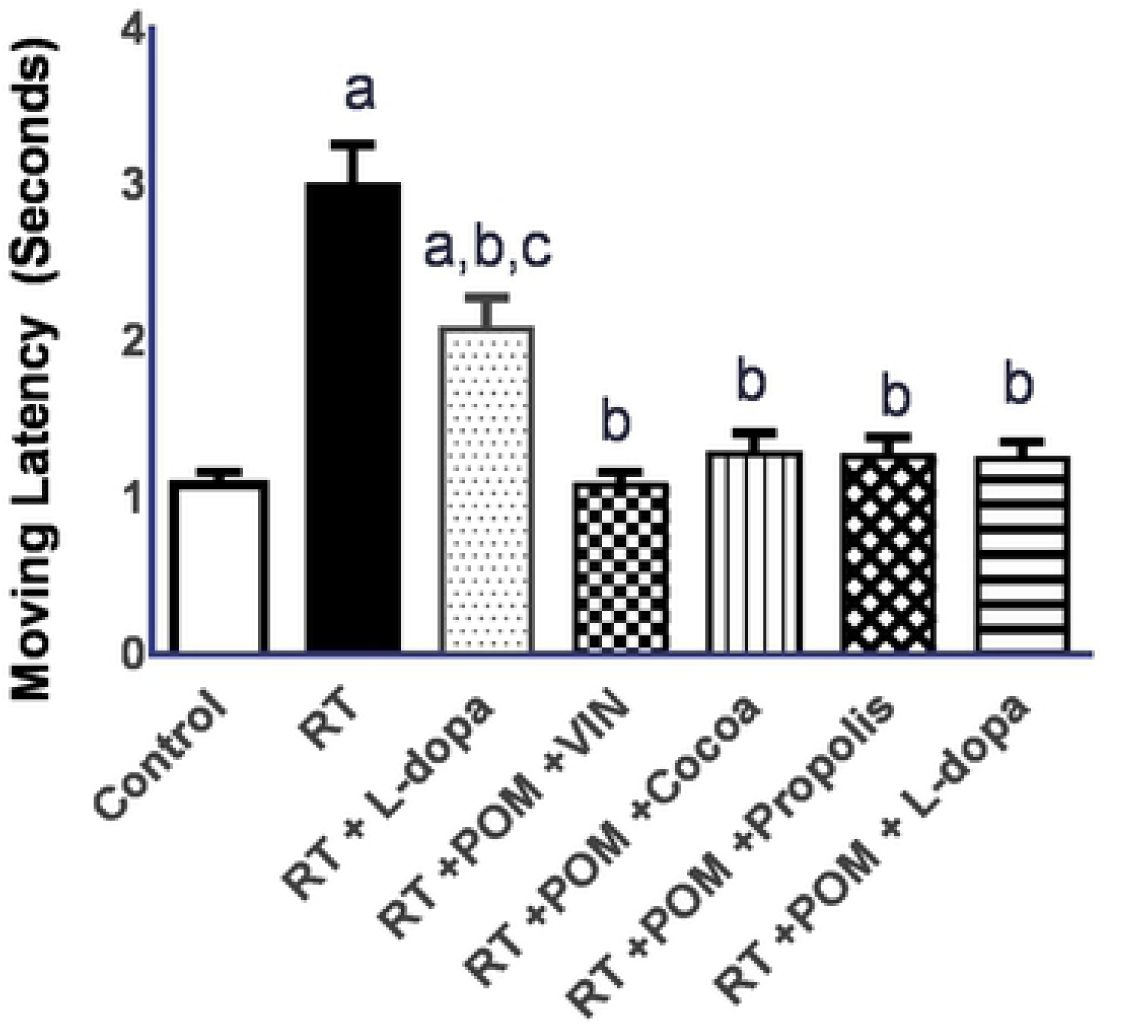
Neurobehavioural Assessment: Moving latency in catalepsy test: a, b, c denote significant difference from control, Rotenone, and Rotenone/L-DOPA groups respectively at P <0.05.

### Effect of Pomegranate together with either of vinpocetine, propolis, cocoa or L-dopa on rotenone-induced changes in swimming latency

Rotenone showed the highest swimming latency, whereas L-dopa or POM alone and/or in combination with VIN, Cocoa and Propolis showed an improvement in comparison to the rotenone group (P<0.05). Swimming latency for the groups: control, rotenone, rotenone and L-dopa, rotenone, POM, and VIN, rotenone, POM, and cocoa, rotenone, POM, and propolis, rotenone, POM and L-dopa were respectively as following: 1.250±0.2739, 6.333±1.506,4.167±0.9309, 2.417±0.3764, 3.167±0.6055, 3.083±0.6646, 3.167±0.5164 seconds) (see figure 5).

**Figure 5:**
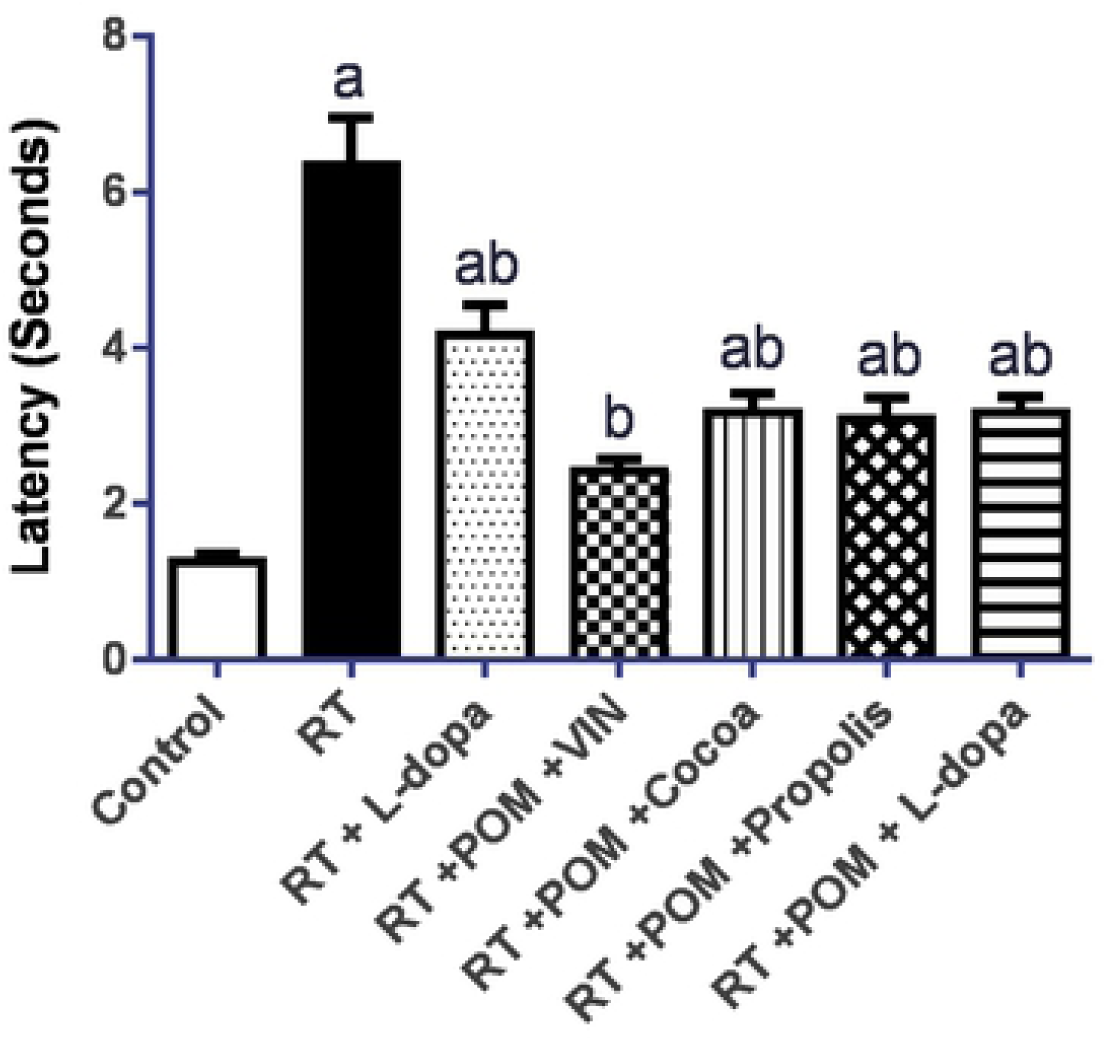
Neurobehavioural assessment: Swimming latency in catalepsy test. a, b denote significant difference from control, and Rotenone groups respectively at P <0.05.

### Effect of pomegranate together with each of vinpocetine, propolis, cocoa or L-dopa on rotenone-induced changes in swimming time

Rotenone showed the highest swimming time, whereas L-dopa or POM alone and/or in combination with VIN, cocoa and propolis showed an improvement in comparison to the rotenone group (P<0.05). Swimming time for the groups: control, rotenone, rotenone and L-dopa, rotenone, POM, and VIN, rotenone, POM, and cocoa, rotenone, POM, and propolis, rotenone, POM and L-dopa were respectively as following: 11.17± 2.714, 20.67±4.967,17.83±3.971, 11.67±1.966, 14.83±2.229, 14.17±1.835, 15.33± 2.251 seconds) (see figure 6).

**Figure 6:**
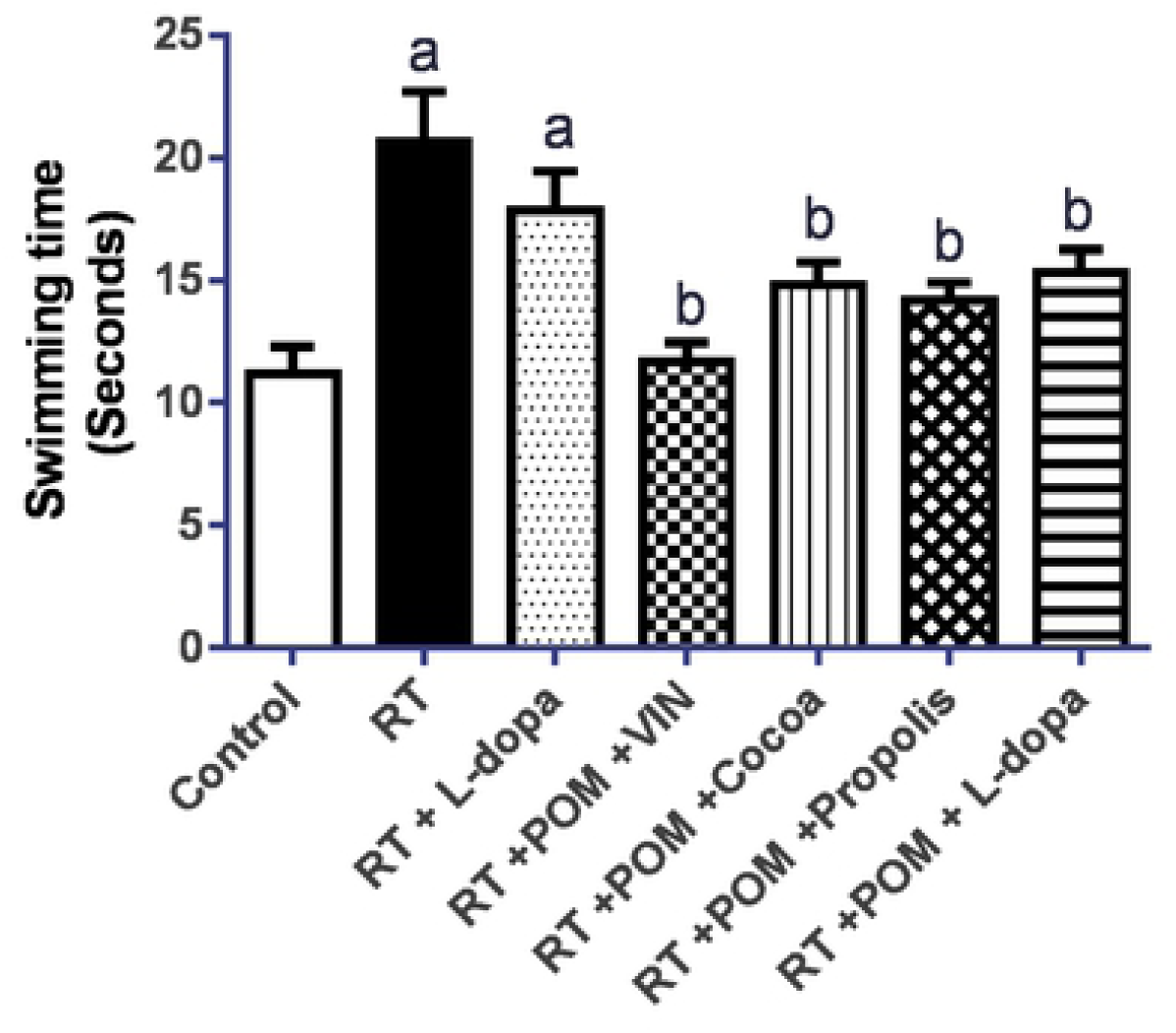
Neurobehavioural assessment: Swimming time in catalepsy test. a, b denote significant difference from control and Rotenone groups respectively at P <0.05.

### Effect of pomegranate together with each of vinpocetine, propolis, cocoa or L-dopa on rotenone-induced changes in swimming score

Rotenone showed the lowest swimming score, whereas L-dopa or POM alone and/or in combination with VIN, cocoa and propolis showed an improvement in comparison to the rotenone and group (P<0.05). For the groups: control, rotenone, rotenone and L-dopa, rotenone, POM, and VIN, rotenone, POM, and cocoa, rotenone, POM, and propolis, rotenone, POM and L-dopa were respectively as following: 3.667±0.5164, 1.833±0.7528, 3.333±0.5164, 3.500±0.5477, 3.167±0.7528, 3.333±0.5164, 3.667±0.5164) (see figure 7).

**Figure 7:**
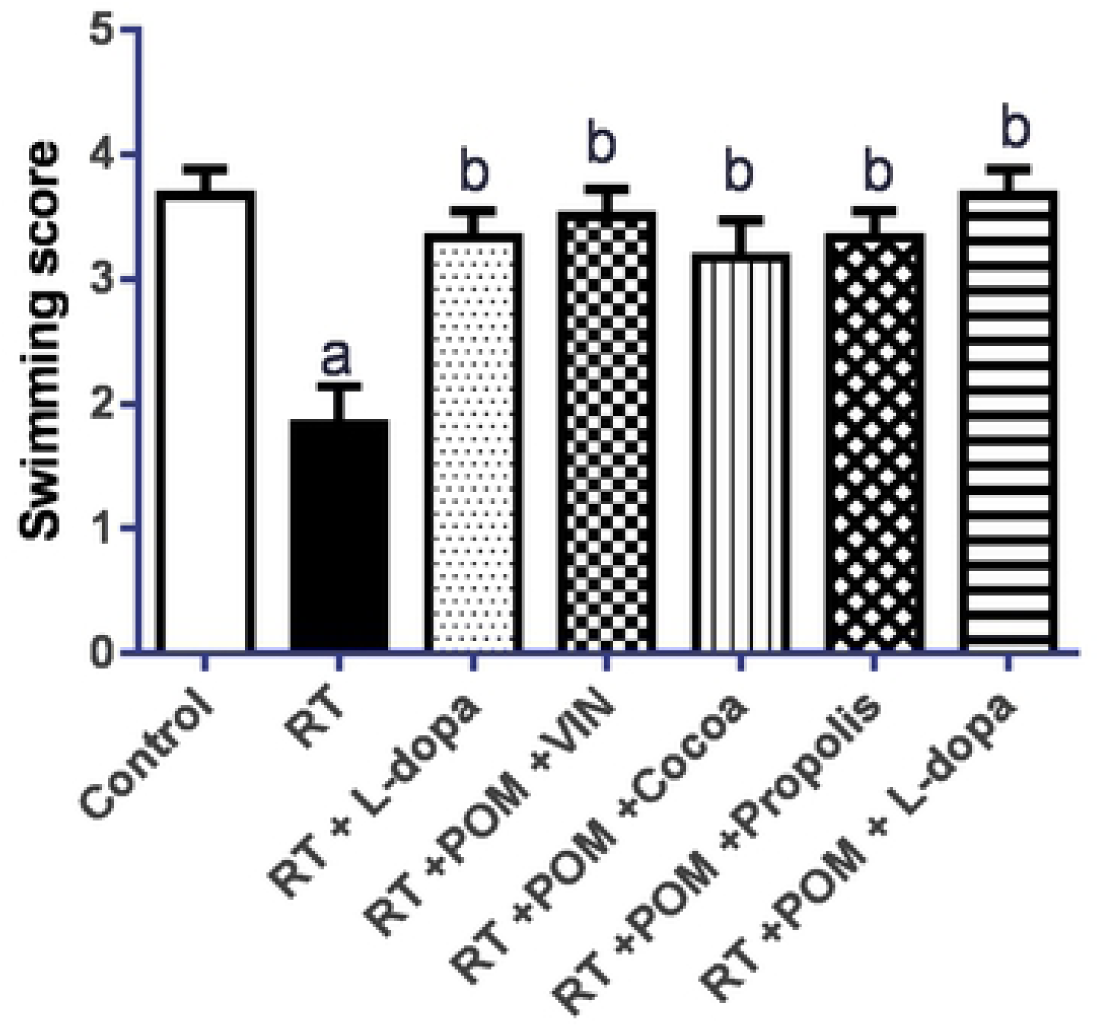
Neurobehavioural assessment: Swimming score in catalepsy test. a, b denote significant difference from control and Rotenone groups respectively at P <0.05.

### Y-maze test

Spatial working memory, a form of short-term memory, can be determined via investigation of spontaneous alternation behavior in a Parkinsonism rat model. Rotenone group showed the lowest percentage of spontaneous alternations, whereas L-dopa or POM alone and/or in combination with VIN, cocoa and propolis showed an improvement in comparison to the rotenone group (P<0.05). The changes in spontaneous alternations for the following groups: control, rotenone, rotenone and L-dopa, rotenone, POM, and VIN, rotenone, POM, and cocoa, rotenone, POM, and propolis, rotenone, POM and L-dopa were respectively as following: 81.17± 2.41, 60.33±1.49, 74.17±3.34, 74.50±2.63, 73.33±1.97, 73.17±2.67, 81.50±2.63) (see figure 8).

**Figure 8:**
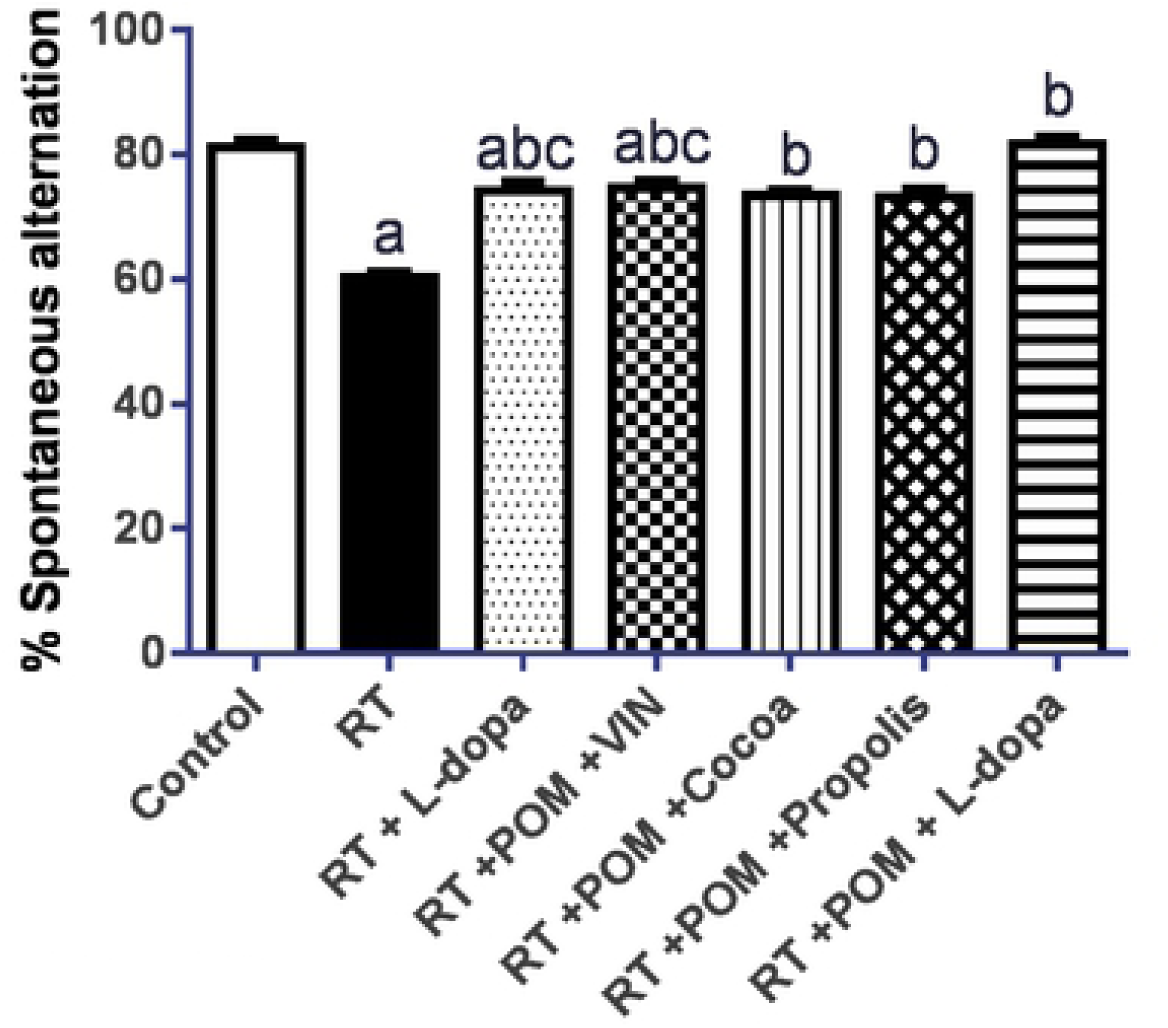
Neurobehavioural assessment: Spontaneous alternations in the Y-maze test. a, b, c denote significantly different from control, Rotenone, and Rotenone/L-DOPA groups respectively at P <0.05.

### The Biochemical efficacies of Pomegranate with different drug Pomegranate, combinations significantly elevated the striatal content of Dopamine, Norepinephrine, and serotonin (5-HT)

Changes in catecholamine levels were measured by ELISA in the striatum (Table 1). Rotenone depleted dopamine, norepinephrine, and serotonin striatal levels. On the other hand, Supplementation of rats with L-dopa or POM alone and/or in combination with VIN, cocoa and propolis showed a significant improvement in the above catecholamine in comparison to the rotenone group (P<0.05).

**Table 1:** Biochemical parameters for all experimental groups.

|  | Control | RT | RT + L-dopa | RT +POM +VIN | RT +POM +Cocoa | RT +POM +Propolis | RT +POM + L-dopa |
| --- | --- | --- | --- | --- | --- | --- | --- |
| **DA** (ng/g) | 72.18±6.0 | 16.68±1.7 | 31.85±2.1 | 45.13±1.9 | 34.42±5.1 | 37.33±3.8 | 62.02±2.4 |
| **NE** (nmol/g) | 574.45±17.6 | 192.88±8.5 | 372.72±5.7 | 398.83±23.0 | 359.04±12.9 | 389.88±21.3 | 475.93±14.3 |
| **5-HT** (ng/g) | 11.18±0.8 | 3.57±0.5 | 4.47±0.7 | 7.45±1.1 | 6.30±1.1 | 6.30±0.7 | 10.50±0.7 |
| **ACHE** (ng/g) | 6.08±0.6 | 31.18±2.5 | 10.03±0.9 | 12.65±1.1 | 17.78±1.3 | 15.40±0.9 | 8.12±0.7 |
| **GABA** | 1.07±0.1 | 8.93±1.1 | 4.62±0.2 | 4.83±0.2 | 5.20±0.7 | 5.29±0.6 | 3.96±0.3 |
| **Glutamate** (ng/ml) | 46.77±1.9 | 11.20±1.0 | 18.18±1.36 | 21.80±1.2 | 17.38±1.4 | 17.63±1.3 | 29.87±1.66 |
| **BDNF** (U/g) | 156.90±5.8 | 59.30±5.3 | 109.13±5.5 | 109.83±4.7 | 108.33±3.5 | 108.45±3.6 | 127.00±3.1 |
| **SOD** (U/g) | 6.66±0.9 | 0.74±0.4 | 2.17±0.3 | 4.96±0.1 | 2.81±0.2 | 2.81±0.2 | 2.58±0.2 |
| **TAC** (μmol/g) | 46.77±1.9 | 11.20±1.0 | 18.18±1.4 | 21.71±1.3 | 17.38±1.4 | 17.38±1.4 | 29.87±1.7 |
| **TNF-α** (pg/g) | 19.08±1.8 | 95.60±1.9 | 47.73±1.6 | 46.43±3.2 | 50.22±2.6 | 50.63±1.6 | 44.17±2.4 |
| **IL1β** (pg/g) | 17.03±1.8 | 115.82±3.2 | 63.40±2.2 | 56.58±5.7 | 67.50±4.9 | 70.22±1.6 | 46.60±2.1 |
| **Caspase3** (ng/ml) | 1.77±0.2 | 18.43±1.5 | 8.82±1.5 | 8.88±0.9 | 11.27±2.1 | 10.78±1.6 | 6.30±0.9 |
| **INOs** (U/mg) | 1.55±0.2 | 43.75±1.9 | 19.85±1.3 | 16.05±2.1 | 18.33±1.3 | 29.87±1.7 | 8.92±1.5 |
| **GSK3B** (fold change) | 1.33±0.3 | 7.90±0.8 | 4.64±0.5 | 4.98±0.3 | 5.45±0.24 | 5.33±0.4 | 3.50±0.3 |

### Pomegranate, vinpocetine, propolis, cocoa or L-dopa decreased the rotenone-induced elevation in the levels of ACHE, GSK3B, and GABA

Rotenone group showed the highest values for ACHE, GSK3B, GABA, whereas L-dopa or POM alone and/or in combination with VIN, cocoa and propolis showed a significant reduction in these parameter compared to the rotenone group (P<0.05) (Table 1).

### Effect of pomegranate, vinpocetine, propolis, cocoa or L-dopa in comparison with L-dopa alone on rotenone-induced changes in Glutamate and BDNF

Rotenone showed the lowest values for Glutamate and BDNF, whereas L-dopa or POM alone and/or in combination with VIN, cocoa and propolis could restore their levels in comparison to the rotenone group (P<0.05) (Table 1).

### Pomegranate, vinpocetine, propolis, cocoa or L-dopa inhibited lipid peroxidation and restored the levels of SOD and TAC

Rotenone group showed the highest values for of MDA, a biomarker for lipid peroxidation, and the lowest antioxidant enzyme level (SOD) and Total antioxidant capacity (TAC), whereas L-dopa or POM alone and/or in combination with VIN, Cocoa and Propolis showed an improvement in comparison to the rotenone group (P<0.05) (Table 1).

### Effect of pomegranate, vinpocetine, propolis, cocoa or L-dopa attenuated the elevated expression of the proinflammatory cytokines TNF-α, IL1β

To examine the possible proinflammatory role of POM and other combinations, we also studied the expression of the cytokine interleukin-1β (IL-1β), tumor necrosis factor-α (TNF-α) in rats exposed to rotenone. In the current study, striatal TNF-α and IL1β levels were increased in the brains of rotenone group as compared to the normal healthy control group, whereas L-dopa or POM alone and/or in combination with VIN, Cocoa and Propolis showed an improvement in comparison to the rotenone group (P<0.05).

### Effect of pomegranate, vinpocetine, propolis, cocoa or L-dopa attenuated the elevated expression of the inflammatory enzyme mediator iNOs

Induction of iNOs is thought to contribute to DA neuron demise. The neurotoxin rotenone increased the expression of iNOS compared to the control group. Co-administration of POM and other combinations in the current study or even L-dopa decreased iNOS compared to the rotenone-alone group. Indeed, Rotenone showed the lowest percentage of iNOS, whereas POM alone and/or in combination with VIN, Cocoa, and Propolis showed an improvement in comparison to the rotenone group (P<0.05) (Table 1).

### Effect of pomegranate, vinpocetine, propolis, cocoa or L-dopa attenuated the elevated expression of Caspase 3

Rotenone induced oxidative damage can lead to caspase 3 activation. In the current study, Rotenone showed an elevation in the level of Caspase 3, whereas POM alone and or in combination with VIN, Cocoa, and Propolis showed a significant decrease in comparison to the rotenone group (P<0.05) (Table 1). Co-administration of POM and other combinations in the current study or even L-dopa decreased Caspase 3 compared to the rotenone-alone group.

### Histopathological study

Figure 9 shows the histopathological analysis conducted for the tissue specimens for the experimental groups employed in the current study showing marked degeneration for the Rotenone group, whereas the rest of the treated groups improved this degeneration**: A. Normal Healthy Control**: No histopathological alterations in the neuronal cells of the cerebral cortex, hippocampus (subiculum, fascia dentate and hilus) and striatum, **B. Rotenone:** Nuclear pyknosis and degeneration in the cerebral cortex are shown in the images, no alterations in the neurons of the subiculum of the hippocampus but nuclear pyknosis in some neurons of the fascia dentate and hilus, multiple focal eosinophilic plagues with nuclear pyknosis and degeneration in some neurons of the striatum, **C. RT + L-Dopa**: Mild congestion in the blood vessels of the striatum and hemorrhage in the meninges, in the cerebellum, **D. RT + POM + VIN**: Intact histological structure of the cerebral cortex, degeneration & nuclear pyknosis in the subiculum of hippocampus while no alteration in the neurons of the fascia dentate and hilus as well as in the striatum, **E. RT + POM +Cocoa**: Nuclear pyknosis and degeneration in the cerebral cortex, no histopathological alterations in the hippocampus (subiculum & fascia dentate) but multiple focal eosinophilic plagues formation with nuclear pyknosis and degenerations in some neurons of the striatum, **F. RT + POM + Propolis**: Nuclear pyknosis and degeneration in the cerebral cortex while few ones in the subiculum in hippocampus had degeneration and nuclear pyknosis, multiple focal eosinophilic plagues formation with nuclear pyknosis and degeneration in some neurons of the striatum, **G. RT + POM + L-Dopa:** No alteration in the cerebral cortex, subiculum and fascia dentate of the hippocampus but nuclear pyknosis and degenerations in some few neurons in the striatum.

**Figure 9:**
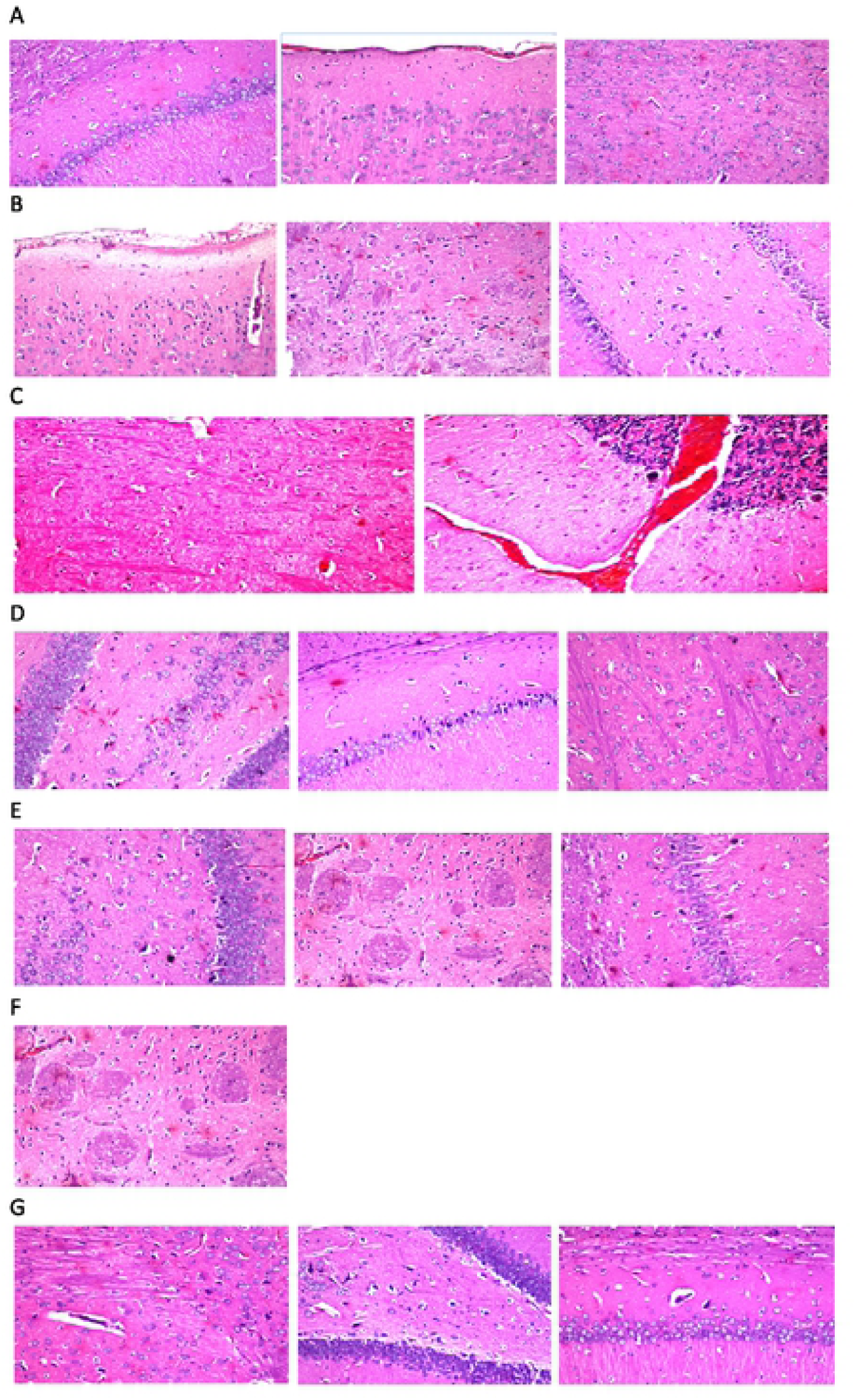
Histopathological investigation: A. Normal Healthy Control. B. Rotenone. C. Rotenone + L-Dopa. D. Rotenone + POM + VIN. E. Rotenone + POM +Cocoa. F. Rotenone + POM + Propolis: G. Rotenone + POM + L-Dopa.

## Discussion

Parkinsonism is a geriatric disorder in which age, genetic and environmental factors are considered as predisposing factors for Parkinsonism. Several lines of evidence indicate oxidative stress and inflammation as important pathogenic factors in this disease. Besides, it was recorded previously that parkinsonism is associated with decreased striatal dopamine levels. Unfortunately, the details of the neurodegenerative cascade of Parkinsonism disease remain unclear. However, it was proposed previously that oxidative damage plays a crucial role in the neuropathology of Parkinsonism. Moreover, Complex I was reported to be impaired in parkinsonism which leads to NADH accumulation resulting in an excessive Reactive Oxygen Species (ROS) production. In agreement with this, recent studies demonstrated an accumulation of MDA in patients with parkinsonism. Increased generation of ROS leads to elevation of the lipid peroxidation level yielding malondialdehyde (MDA) which can be considered as a biomarker for oxidative stress. Also, other factors observed in Parkinsonism pathology are glutathione (GSH) depletion, enhanced superoxide activity, and cellular apoptosis [11,12].

L-dopa was always considered as the best treatment for parkinsonism despite its long-term treatment side effects which may include further addition to the free radical oxidative load produced via normal metabolism processes. This may lead to the progression of the disease. Therefore, L-dopa is only limited to the relief of Parkinsonism symptoms. Therefore, developing new agents that decrease the rate of Parkinsonism progression is crucial [13,14]. Since the key factor to relief, the oxidative stress seen in Parkinsonism is to repair the damage due to free radicals via antioxidants which might have a preventive action against free radical-mediated tissue destruction, a combination of Pomegranate, propolis, vinpocetine and cocoa will be investigated in the current study.

Rotenone is a pesticide of lipophilic nature and hence can cross biological membranes. Rotenone is known to mimic the neurochemical and behavioral features of Parkinsonism in rats. Indeed, rotenone impaired behavioral and biochemical profile of the rats under investigation in the current study. Examples for such features are reduced mobility and flexed posture [15]. It was proposed previously that this is because rotenone damages neurons via oxidative mechanisms and hence increases lipid peroxidation. Rotenone-treated rats have previously demonstrated elevated levels of TNF-α. The released TNF-α may be responsible for neuronal degeneration in parkinsonism disease.

POM with different combinations significantly reversed the deficits observed with rotenone. Pomegranate, Punica granatum L., is a fruit consumed fresh or as a juice [16]. Pomegranate possesses a well-known health-promoting effect. Pomegranate fruits are rich in polyphenols. Indeed, 23 phenolic compounds were identified including cyanidin-3,5-O-diglucoside and pelargonidin-3,5-O-diglucoside. It was previously indicated that polyphenols attenuate neuronal death in animal models of neurodegeneration. This can be explained based on their possession to antioxidant and anti-inflammatory actions [16,17].

Vinpocetine (VIN) is a synthetic derivative of the alkaloid vincamine. It is a broad-spectrum antioxidant and neuroprotective agent. It was reported previously to have a therapeutic effect in the treatment of some cerebrovascular diseases. It was previously demonstrated that vinpocetine inhibited oxidative stress, memory impairment, and neuroinflammation via enhancement of the antioxidant defense system and inhibition of neuroinflammatory cytokines. The neuroprotective effect of VIN may be due to modulation of monoamines, antioxidant, anti-inflammatory, and anti-apoptotic activities [10]. In addition to that, alteration of the rheological properties of red blood cells is a unique mechanism of VIN. This unique mechanism enables the penetration of the small vessels of the cerebro-microvasculature by blood cells to deliver nutrients improving neurocognitive function. Findings from a previous study demonstrated that vinpocetine improved the symptoms of parkinsonism via improvement of the antioxidant defense system and inhibition of neuroinflammatory mediators. Moreover, VIN significantly reduced MDA and increased GSH levels as compared to the control positive group [5,18].

It was reported previously that propolis with the caffeic acid phenethyl ester, as the main active component, attenuated dopaminergic neurodegeneration in a mouse model of Parkinsonism disease [19]. Propolis and cocoa were shown previously to possess anti-inflammatory as well as antioxidant properties [19,20].

In the current study, different levels of biochemical markers were assessed in addition to histopathological examinations of different brain regions were also determined.

Brain-derived neurotrophic factor (BDNF) is secreted in the mammalian central nervous system. Diminished BDNF expression level in the substantia nigra in the animal can mimic the symptoms of Parkinsonism. In the present study, POM with other combinations elevated the decreased level of BDNF following the injection of rotenone [21]. In addition to that, the elevated glycogen synthase kinase (GSK 3B) level, which was implicated in the pathogenesis of several neurodegenerative diseases such as Alzheimer’s disease, was reported to decrease following administration of POM with other combinations [22].

The authors of this study were able to show that combinations of POM with each of VIN, Propolis or Cocoa have beneficial effects against the development of Parkinsonism.

## Acknowledgement

We acknowledge Alexandria university and Azhar university for funding the current research study

## References

[1] E. Botsford, J. George, E.E. Buckley, Parkinson’s Disease and Metal Storage Disorders: A Systematic Review., Brain Sci. 8 (2018). doi: 10.3390/brainsci8110194.

[2] W.C. Ballance, E.C. Qin, H.J. Chung, M.U. Gillette, H. Kong, Reactive oxygen species-responsive drug delivery systems for the treatment of neurodegenerative diseases., Biomaterials. 217 (2019) 119292. doi: 10.1016/j.biomaterials.2019.119292.

[3] N. Bhide, D. Lindenbach, M.A. Surrena, A.A. Goldenberg, C. Bishop, S.P. Berger, M.A. Paquette, The effects of BMY-14802 against L-DOPA- and dopamine agonist-induced dyskinesia in the hemiparkinsonian rat., Psychopharmacology (Berl). 227 (2013) 533–544. doi: 10.1007/s00213-013-3001-4.

[4] G. Kaur, Z. Jabbar, M. Athar, M.S. Alam, Punica granatum (pomegranate) flower extract possesses potent antioxidant activity and abrogates Fe-NTA induced hepatotoxicity in mice., Food Chem. Toxicol. 44 (2006) 984–993. doi: 10.1016/j.fct.2005.12.001.

[5] I.O. Ishola, A.A. Akinyede, T.P. Adeluwa, C. Micah, Novel action of vinpocetine in the prevention of paraquat-induced parkinsonism in mice: involvement of oxidative stress and neuroinflammation., Metab. Brain Dis. 33 (2018) 1493–1500. doi: 10.1007/s11011-018-0256-9.

[6] M. Messaoudi, J.-F. Bisson, A. Nejdi, P. Rozan, H. Javelot, Antidepressant-like effects of a cocoa polyphenolic extract in Wistar-Unilever rats., Nutr. Neurosci. 11 (2008) 269–276. doi: 10.1179/147683008X344165.

[7] U.Z. Usman, A.B.A. Bakar, M. Mohamed, Propolis improves pregnancy outcomes and placental oxidative stress status in streptozotocin-induced diabetic rats., BMC Complement. Altern. Med. 18 (2018) 324. doi: 10.1186/s12906-018-2391-6.

[8] L.H. Morais, B.J. Martynhak, R. Santiago, T.T. Takahashi, D. Ariza, J.K. Barbiero, R. Andreatini, M.A.B.F. Vital, M.M.S. Lima, Characterization of motor, depressive-like and neurochemical alterations induced by a short-term rotenone administration, Pharmacol. Reports. 64 (2012) 1081–1090. doi: 10.1016/S1734-1140(12)70905-2.

[9] R.-X. Xu, N. Grigoryev, T.-L. Li, H.-S. Bian, R. Zhang, X.-Y. Liu, Development of hexagonal maze procedure for evaluating memory in rat., Biomed. Reports. 1 (2013) 134–138. doi: 10.3892/br.2012.16.

[10] S.A. Zaitone, D.M. Abo-Elmatty, S.M. Elshazly, Piracetam and vinpocetine ameliorate rotenone-induced Parkinsonism in rats., Indian J. Pharmacol. 44 (2012) 774–779. doi: 10.4103/0253-7613.103300.

[11] T. Wichmann, Changing views of the pathophysiology of Parkinsonism., Mov. Disord. (2019). doi: 10.1002/mds.27741.

[12] J. Silva, C. Alves, R. Freitas, A. Martins, S. Pinteus, J. Ribeiro, H. Gaspar, A. Alfonso, R. Pedrosa, Antioxidant and Neuroprotective Potential of the Brown Seaweed Bifurcaria bifurcata in an in vitro Parkinson’s Disease Model., Mar. Drugs. 17 (2019). doi: 10.3390/md17020085.

[13] M.J. Kelly, M.A. Lawton, F. Baig, C. Ruffmann, T.R. Barber, C. Lo, J.C. Klein, Y. Ben-Shlomo, M.T. Hu, Predictors of motor complications in early Parkinson’s disease: A prospective cohort study., Mov. Disord. (2019). doi: 10.1002/mds.27783.

[14] N. Malek, S. Kanavou, M.A. Lawton, V. Pitz, K.A. Grosset, N. Bajaj, R.A. Barker, Y. Ben-Shlomo, D.J. Burn, T. Foltynie, J. Hardy, N.M. Williams, N. Wood, H.R. Morris, D.G. Grosset, L-dopa responsiveness in early Parkinson’s disease is associated with the rate of motor progression., Parkinsonism Relat. Disord. (2019). doi: 10.1016/j.parkreldis.2019.05.022.

[15] K.R. Rekha, R. Inmozhi Sivakamasundari, Geraniol Protects Against the Protein and Oxidative Stress Induced by Rotenone in an In Vitro Model of Parkinson’s Disease., Neurochem. Res. 43 (2018) 1947–1962. doi: 10.1007/s11064-018-2617-5.

[16] V. Tapias, J.R. Cannon, J.T. Greenamyre, Pomegranate juice exacerbates oxidative stress and nigrostriatal degeneration in Parkinson’s disease., Neurobiol. Aging. 35 (2014) 1162–1176. doi: 10.1016/j.neurobiolaging.2013.10.077.

[17] M.K. Mazumder, S. Choudhury, A. Borah, An in silico investigation on the inhibitory potential of the constituents of Pomegranate juice on antioxidant defense mechanism: Relevance to neurodegenerative diseases., IBRO Reports. 6 (2019) 153–159. doi: 10.1016/j.ibror.2019.05.003.

[18] R.I. Nadeem, H.I. Ahmed, B.M. El-Sayeh, Protective effect of vinpocetine against neurotoxicity of manganese in adult male rats., Naunyn. Schmiedebergs. Arch. Pharmacol. 391 (2018) 729–742. doi: 10.1007/s00210-018-1498-0.

[19] S.A. Zaitone, E. Ahmed, N.M. Elsherbiny, E.T. Mehanna, M.K. El-Kherbetawy, M.H. ElSayed, D.M. Alshareef, Y.M. Moustafa, Caffeic acid improves locomotor activity and lessens inflammatory burden in a mouse model of rotenone-induced nigral neurodegeneration: Relevance to Parkinson’s disease therapy., Pharmacol. Rep. 71 (2019) 32–41. doi: 10.1016/j.pharep.2018.08.004.

[20] I. Cova, V. Leta, C. Mariani, L. Pantoni, S. Pomati, Exploring cocoa properties: is theobromine a cognitive modulator?, Psychopharmacology (Berl). 236 (2019) 561–572. doi: 10.1007/s00213-019-5172-0.

[21] C.M. Costa, G.L. de Oliveira, A.C.S. Fonseca, R. de C. Lana, J.C. Polese, A.P. Pernambuco, Levels of cortisol and neurotrophic factor brain-derived in Parkinson’s disease., Neurosci. Lett. 708 (2019) 134359. doi: 10.1016/j.neulet.2019.134359.

[22] A. Petit-Paitel, [GSK-3beta: a central kinase for neurodegenerative diseases?]., Med. Sci. (Paris). 26 (2010) 516–521. doi: 10.1051/medsci/2010265516.

